# Targeting ribosome biogenesis sensitizes PARPi-resistant triple-negative breast cancer

**DOI:** 10.64898/2026.10.08.757577

**Authors:** Mallory I. Frederick, Nicolas Pouderous, Kenan Aloss, Qi Yang, Gerardo Ferbeyre, Saima Hassan

**Author notes:** Corresponding author: Saima Hassan, MD, PhD, FRCSC, CRCHUM, 900 St. Denis Street, Pavilion R, R10.422, Montréal, Québec, H2X 0A9. Authors contributed equally (Mallory I. Frederick, Nicolas Pouderous).

## Abstract

Poly (ADP-ribose) polymerase inhibitors (PARPi) are the only targeted treatment available for early triple-negative breast cancer (TNBC), an aggressive subtype with limited treatment options. While PARPi are effective in patients with BRCA mutations, resistance to these inhibitors can arise. We previously developed BRCA-mutant cell lines, HCC1395, with acquired resistance to talazoparib and olaparib and have now developed PARPi-resistant MX1 cell lines in an analogous manner. Ribosomal pathways were upregulated in 3/4 cell lines with acquired PARPi resistance. The resistant cell lines showed high and low basal nucleolar stress based on nucleolar morphology and nucleophosmin delocalization from nucleolus to nucleoplasm. Treatment of these cell lines with the combination of olaparib and a ribosome biogenesis inhibitor, BMH-21, demonstrated synergy in the PARPi-resistant cell lines, but not in the parental cells. Interestingly, HCC1395 cells, which had lower baseline nucleolar stress, showed greater nucleophosmin delocalization with combination treatment, than MX1 cells, which had higher baseline nucleolar stress. All four cell lines demonstrated sustained DNA damage with the combination. We extended our investigation to 3 BRCA wild-type TNBC cell lines, BT549, Hs578T, and MDAMB231 and identified synergy in the context of intrinsic PARPi resistance. In MDAMB231 cells, combining olaparib with BMH-21 induced a senescence-like phenotype, increasing γH2AX-positive cells by up to 7.3-fold, senescence-associated β-galactosidase (SA-β-gal) staining by 5.6-fold, and p21 by 3.6-fold. These results were mirrored with a second ribosome biogenesis inhibitor, CX-5461. Finally, we identified distinct patterns of ribosomal pathway activity in a cohort of patients with treatment-resistant TNBC, and found correlations between ribosomal pathways with MYC, proliferation, DNA damage, and senescence. Together, these data identify ribosome biogenesis as a potential therapeutic vulnerability in PARPi-resistant TNBC and support its targeting in combination with PARPi, while demonstrating a broader for ribosome biogenesis in treatment-resistant TNBC.

## Introduction

Breast cancer accounts for 24.3% of cancers worldwide and is the leading cause of cancer deaths in women^1^. Among breast cancers, triple-negative breast cancer (TNBC) is the most aggressive subtype, characterized by a lack of overexpression of the estrogen and progesterone receptors (ER/PR) and the human epidermal growth factor receptor 2 (HER2). TNBC patients often receive multiple chemotherapy agents and have limited therapeutic options. The only targeted therapies currently available for early TNBC patients are poly (ADP-ribose) (PAR) polymerase (PARP) inhibitors (PARPi), having demonstrated an improvement in overall survival in breast cancer patients with germline mutations in BRCA1/2^2^.

PARPi, including olaparib and talazoparib, target PARP-1/2 enzymes, which are critical sensors of DNA-strand breaks^3^. Active PARP-1 is known to auto-PARylate, which signals the recruitment of DNA repair proteins to the damage site^4,5^. By inhibiting the catalytic activity of PARP-1/2, PARPi block the formation of PAR polymers, and impair the recruitment of DNA repair proteins. In the context of BRCA1/2-deficient cells, PARPi lead to an accumulation of double-strand breaks, resulting in genomic instability and cell death^6^. PARPi can also act by PARP-DNA trapping, wherein the PARP-1 enzyme is trapped at the site of the DNA damage, preventing both repair and DNA replication^7^. Thus, PARPi cytotoxicity reflects both impared DNA repair and disruption of DNA replication, with PARP-DNA trapping providing an important link between these processes^8^.

Beyond BRCA-mutant disease, PARPi are gaining potential for patients exhibiting BRCAness, characterized by deficiencies causing similar phenotypes to *BRCA1/2* mutations, while lacking the actual mutation^9,10^. BRCAness can arise in patients due to mutations in HR-associated genes, epigenetic mechanisms that impair HR function, copy number alterations indicative of genomic scarring, or mutational signatures^11–13^. Indeed, olaparib was found to have approximately 60% efficacy in a small cohort of BRCA wild-type patients, implicating other pathways in PARPi response. Previously, we demonstrated synergy between talazoparib with LNT1, which targets Flap endonuclease 1 (FEN1) and exonuclease 1 (EXO1), in BRCA wild-type TNBC cell lines and a BRCA2-mutant cell line with acquired resistance to olaparib, HCC1395-OlaR^14^. Interestingly, we identified ribosomal pathways as a common feature of PARPi-resistant TNBC cell lines, suggesting that ribosome biogenesis could be exploited for future therapeutics.

The nucleolus is the primary site of ribosomal RNA (rRNA) transcription and processing in the nucleus. PARP-1, FEN1, and EXO1, in addition to being nuclear proteins, localize to the nucleolus. Indeed, it has been reported that up to 40% of PARP-1 localizes to the nucleolus^15^. Importantly, there is increasing evidence of crosstalk between DNA repair and ribosome biogenesis pathways, with DNA repair factors contributing to the maintenance of nucleolar function and rDNA integrity^16^. When pre-rRNA transcription is disrupted, cells experience nucleolar stress, which is characterized by the de-localization of nucleophosmin (NPM1) from the nucleolus to the nucleoplasm, among other markers^17^. Previous work also linked nucleolar stress to cellular senescence^18^, a stable cell-cycle arrest that can also be induced by DNA damage^19^. These links between ribosome biogenesis, DNA repair, and cellular senescence may be particularly relevant in cancer, where replication stress may interfere with ribosome biogenesis^20^ thus increasing the need for PARP-1 activity in the nucleolus.

Aberrant hyperactivation of ribosome biogenesis has been recognized as a hallmark of cancer associated with poor prognosis^21^. Cancer cells have significantly increased ribosome biogenesis and are therefore more susceptible to treatments targeting this process^21–24^. Ribosomal RNA modifying proteins are an integral component of ribosome biogenesis that have recently been shown to be enriched in TNBCs, correlating with increasing genomic instability, elevated tumor mutational burden and poor outcomes^25^.

Here, we show that inhibition of ribosome biogenesis by an RNA polymerase I (Pol I) inhibitor, BMH-21, in combination with the PARPi olaparib, leads to synergistic effects in TNBC cell lines with acquired and innate PARPi resistance. Amongst cell lines with acquired resistance to PARPi, we observe heterogeneity in baseline characteristics, supporting distinct mechanisms of synergy with BMH-21 and olaparib. The combination of BMH-21 and olaparib increased nucleolar stress, DNA damage, and senescence. These data identify ribosome biogenesis as a therapeutic vulnerability and support its targeting in combination with PARPi in PARPi-resistant and BRCA wild-type TNBC.

## Results

### Targeting ribosome biogenesis enhances olaparib sensitivity in PARPi-resistant TNBC cell lines

To determine whether ribosome function is associated with PARPi resistance, we investigated transcriptional reprogramming in TNBC cell lines with acquired resistance to PARPi. We used the TNBC cell line MX1, which is BRCA2-mutant and BRCA1-deficient^26,27^, to develop cell lines with acquired resistance to olaparib (MX1-OlaR) and talazoparib (MX1-TalaR), at similar levels to our previously developed HCC1395-OlaR and HCC1395-TalaR cell lines^14^ (Figure 1A,B). RNAseq and pre-ranked Gene Set Enrichment Analysis (GSEA) was performed in each resistant cell line and compared with its parental counterpart^14^, using the Hallmark, KEGG, and curated ribosome-related pathways. Three of the four resistant cell lines demonstrated an upregulation of the KEGG_Ribosome pathway, which includes many structural components of the large and small ribosomal subunits (Figure 1C). Interestingly, MX1-TalaR also showed an upregulation of pathways involved in ribosome biogenesis, ribosome assembly, rRNA processing, large subunit ribosome biogenesis and Myc targets. In contrast, MX1-OlaR demonstrated an upregulation of inflammatory-like signalling including cytokine, and cytosolic DNA sensing. Together, these results highlight that upregulation of ribosome-related pathways as a prominent feature in cell lines with acquired PARPi resistance.

**Figure 1.**
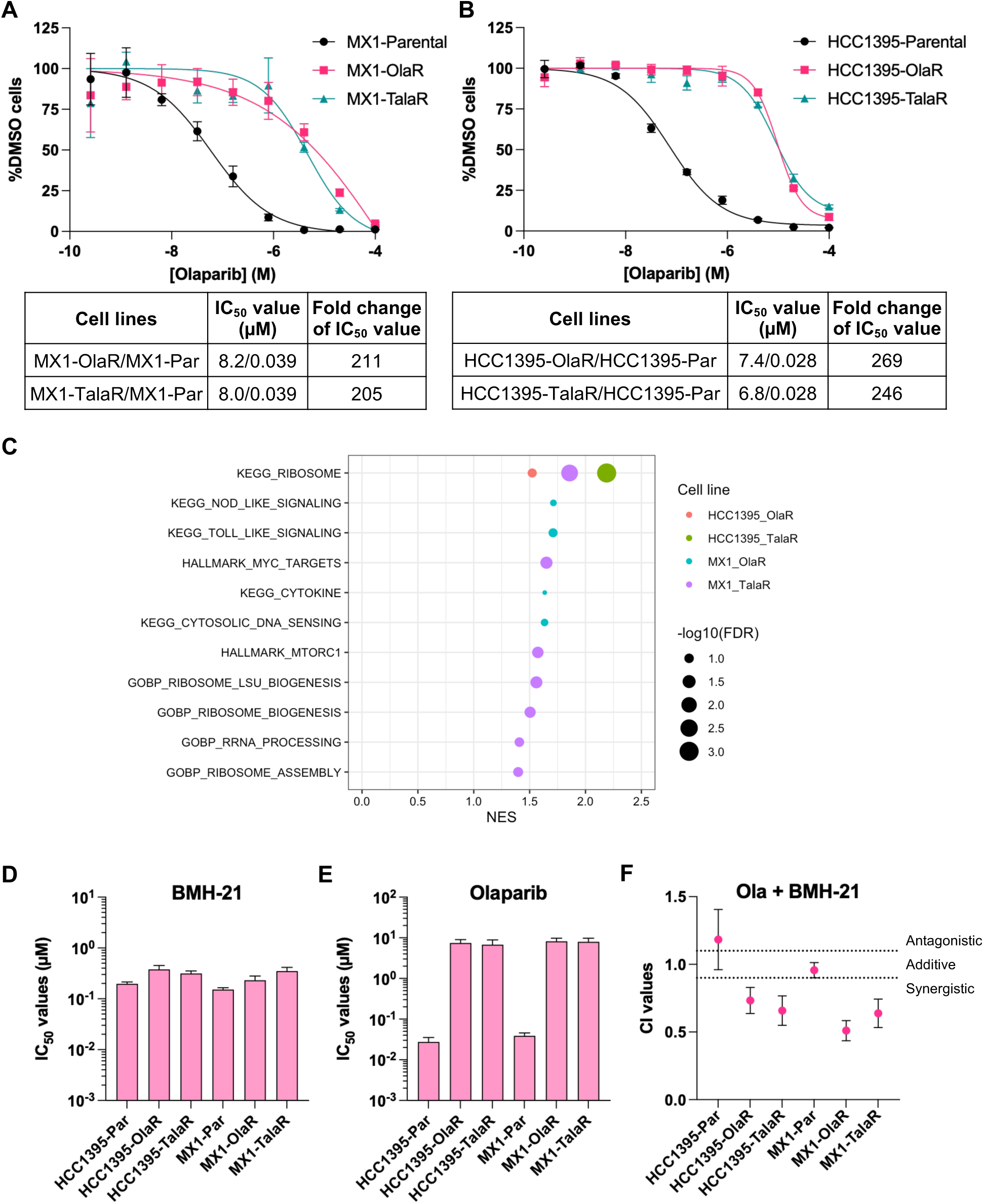
A Pol I inhibitor, BMH-21, induces sensitivity to olaparib in TNBC cells with acquired PARPi resistance. **(A)** MX1 and **(B)** HCC1395 cell lines with acquired resistance to PARPi. IC50 dose-response curves for Parental (Par, black circles), olaparib resistant (OlaR, pink squares), and talazoparib resistant (TalaR, teal triangles) treated with olaparib in a 10-day assay. IC50 values for resistant and parental cell lines and ratio relative to parentals are indicated below. **(C)** Dot plot of pre-ranked GSEA comparing HCC1395/MX1 cell lines with resistance to olaparib or talazoparib in comparison to parental cell lines. Selected significant pathways are shown, with p<0.05, and FDR <0.25. IC50 values for **(D)** BMH-21 and **(E)** olaparib in parental and acquired resistant cell lines. **(F)** Combination index values for olaparib and BMH-21 reported at an FA of 0.5.

The upregulation of ribosome-related pathways in PARPi-resistant cells led us to investigate whether ribosome biogenesis can be therapeutically targeted to enhance PARPi response. Using the Chou and Talalay method^28^, we measured the impact of the combination of olaparib and the Pol I inhibitor BMH-21 in parental and PARPi-resistant HCC1395 and MX1 cell lines in 10-day chemosensitivity assays. The combination demonstrated synergy (Combination index [CI] 0.51 - 0.73, at Fa = 0.5) in all four of our acquired PARPi-resistant cell lines, but not in their respective parental cell lines (Figure 1D-F).

### PARPi resistance alters nucleolar morphology and function

We first evaluated baseline nucleolar morphology of parental, OlaR, and TalaR cell lines derived from HCC1395 and MX1 cells. Using high-content imaging of DAPI-stained cells combined with immunofluorescence of RSL1D1, a nucleolar protein, we measured the size and number of nucleoli (Figure 2A-D). Interestingly, PARPi-resistant HCC1395 and MX1 cell lines showed distinct morphological changes. In HCC1395 cells, the nucleolar area decreased modestly by 9.6-13.1% in OlaR and TalaR cells, with a concomitant 15.6% decrease in the number of nucleoli per cell in TalaR, but not OlaR cells. In contrast, the nucleolar area of MX1-OlaR and MX1-TalaR cells increased by an average of 20.8% relative to MX1-Par cells, with the number of nucleoli increased by 51.7-67.0% in the resistant cells (Figure 2C,D).

**Figure 2.**
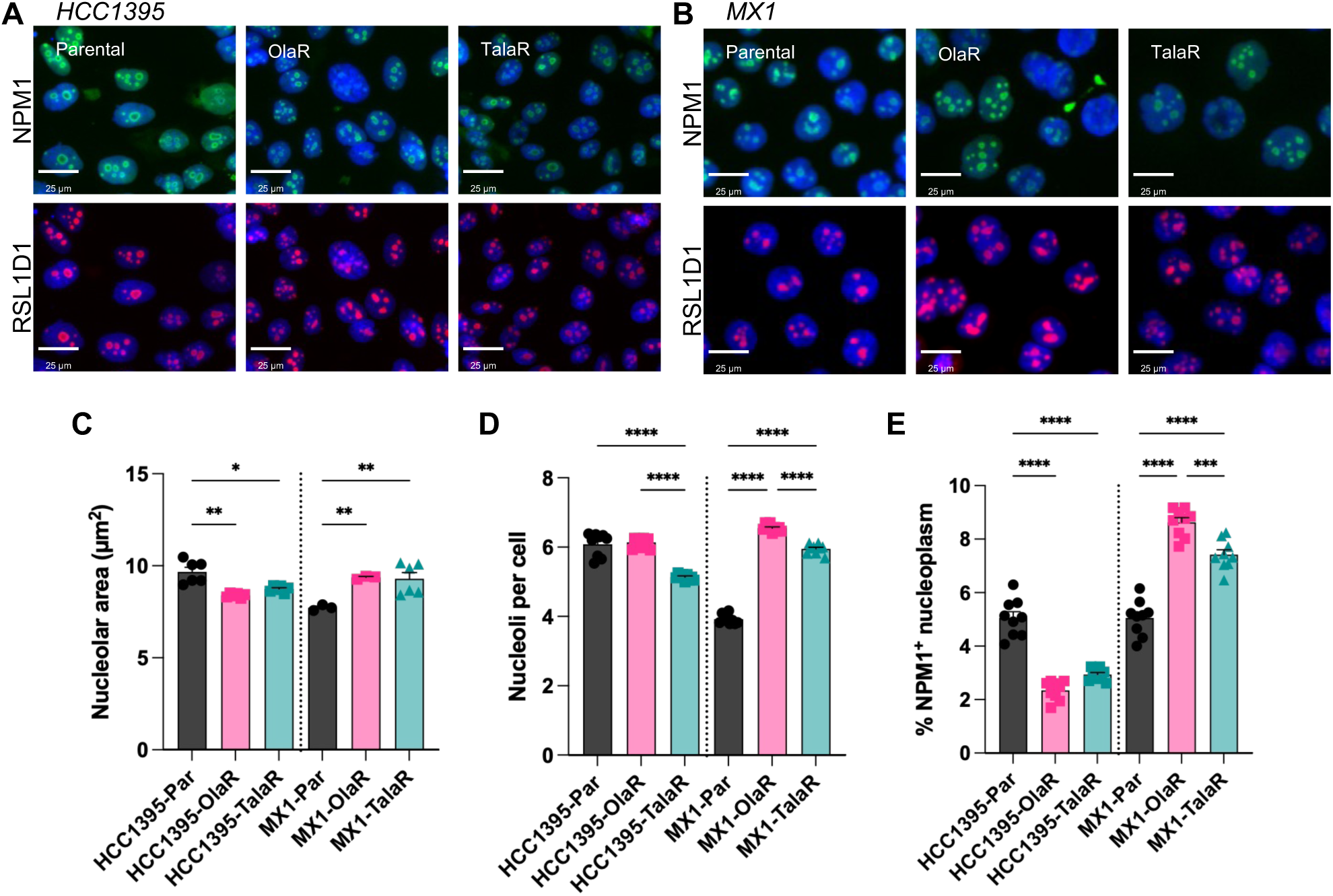
Nucleolar morphology is altered in acquired PARPi resistance. Representative immunofluorescence images of **(A)** HCC1395 and **(B)** MX1 cell lines with acquired PARPi resistance. Images include NPM1 (green) and RSL1D1 (red), with nuclear DAPI counterstaining (blue). Scale bars indicate 25 µm. Quantification of **(C)** nucleolar area, **(D)** number of nucleoli, and **(E)** NPM1 delocalization in HCC1395 and MX1 parental and acquired PARPi-resistant cell lines. Bar graphs indicate mean ± SEM. Two-way ANOVA followed by Tukey’s multiple comparisons was performed to test statistical significance within each starting cell line, with P < 0.0001****, P <0.001***, P < 0.01**, P < 0.05*.

We next used immunofluorescence to compare nucleophosmin (NPM1) in acquired and parental resistance cells (Figure 2A,B). NPM1 is a marker of nuceolar stress that redistributes to the nucleoplasm upon nucleolar disruption^29,30^. Consistent with the changes in nucleolar morphology, HCC1395-OlaR and HCC1395-TalaR cells showed significantly less NPM1 delocalization, with 41.8-53.5% reductions in the proportion of cells exhibiting NPM1 redistribution. In contrast, the proportion of cells with NPM1 redistribution increased by 70.6% and 46.7% in MX1-OlaR and MX1-TalaR cells, respectively (Figure 2E). Together, the nucleolar morphology and NPM1 localization results revealed lower basal nucleolar stress in HCC1395-resistant cells and higher basal nucleolar stress in MX1-resistant cells.

### Combining BMH-21 and olaparib increases nucleolar stress and DNA damage

Given the differences in basal nucleolar stress across the resistant cell lines, we next examined the treatment effects of BMH-21 and olaparib on nucleolar stress and DNA damage. While the treatment-related effects on nucleolar morphology were not as substantive as baseline changes, we observed important trends. The combination increased nucleolar area by 7.5-22.2% in HCC1395 resistant cells, which was comparable to 13.7% increase observed in the parental cells (Figure S1A-C). In MX1 cells, the combination resulted in an increase in nucleolar area by 13.6% with a decrease in the number of nucleoli by 4.4% in resistant cells, while the parental cells demonstrated a 12.7% decrease in nucleolar area associated with a 21.2% increase in number of nucleoli. (Figure S1D-F). The opposing changes in nucleolar area and nucleolar number suggest distinct patterns of treatment-induced nucleolar remodeling in parental and PARPi-resistant MX1 cells, which was not observed with HCC1395 cells.

We next assessed treatment-induced changes in NPM1 localization. The combination of BMH-21 and olaparib increased the proportion of cells with NPM1 redistribution by 4.5-8.1-fold, compared to 3.2-fold in HCC1395-Par. The combination resulted in a 2.5-fold increase NPM1 relocalization MX1-TalaR cells, which was statistically significantly higher than 1.7-1.8-fold increase observed in the MX1-OlaR or MX1-Par cells (Figure 3A,B). While BMH-21 alone contributed to NPM1 relocalization in the HCC1395-OlaR cell line, no such effect was identified in the remaining cell lines. Thus, the combination induced NPM1 relocalization in both HCC1395 and MX1 cells, with the greatest effects observed in HCC1395-resistant and MX1-TalaR cells.

**Figure 3.**
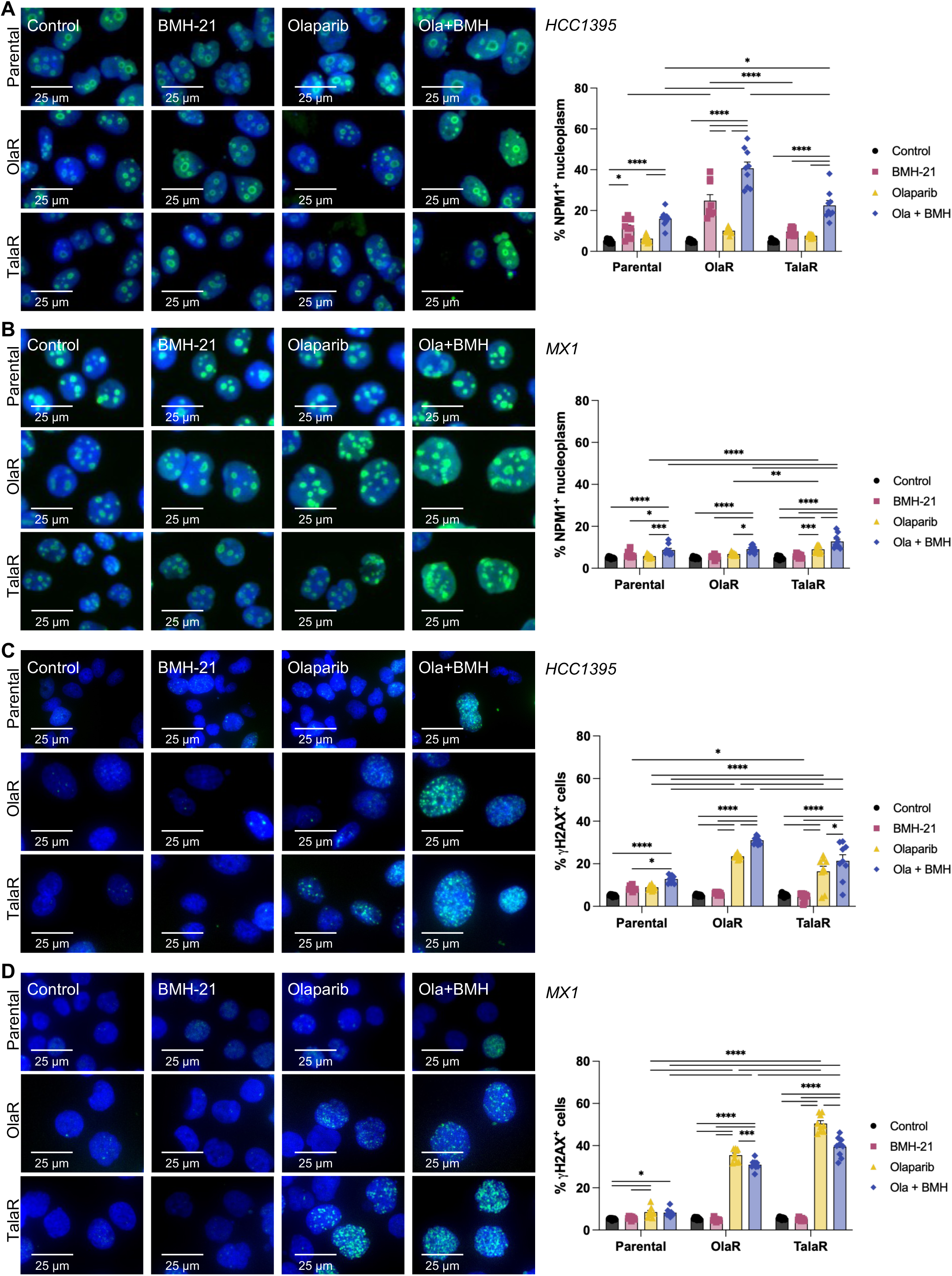
Combined BMH-21 and olaparib induce nucleolar stress and DNA damage in cell lines with acquired PARPi resistance. Representative images and quantification of **(A,B)** NPM1 delocalization or **(C,D)** γH2AX foci formation in **(A,C)** HCC1395 and **(B,D)** MX1 parental, olaparib-resistant, and talazoparib-resistant cell lines treated with DMSO control (black circles), BMH-21 (pink squares), olaparib (yellow triangles), or olaparib combined with BMH-21 (blue diamonds). Images include NPM1 (green) and γH2AX (red), with nuclear DAPI counterstaining (blue). Scale bars indicate 25 µm. Bar graphs of percentage of NPM1 delocalization-positive or γH2AX-positive cells indicate mean ± SEM. Two-way ANOVA followed by Tukey’s multiple comparisons was performed to test statistical significance within each cell line, with P < 0.0001****, P <0.001***, P < 0.01**, P < 0.05*.

Next, we investigated the impact of treatment on the accumulation of DNA damage. The combination of olaparib with BMH-21 resulted in a 4.0-6.0-fold increase in the proportion of γH2AX-positive HCC1395-OlaR/TalaR cells, compared with a 2.5-fold increase in HCC1395-Par cells. In MX1, the combination resulted in 5.7-9.1-fold increase in γH2AX-positive cells in comparison to the 1.6-fold increase in the parental counterpart (Figure 3C,D). Of note, olaparib-induced DNA damage played an important role to the combination effect, however statistically significant differences between the combination and olaparib were still observed in HCC1395. Therefore, all PARPi-resistant cell lines demonstrated increased DNA damage in response to the combination treatment, with a more pronounced effect observed in MX1 cells.

Finally, we investigated whether the olaparib and BMH-21 combination treatment was associated with the induction of cyclin-dependent kinase inhibitor p21, a key mediator of therapy-induced senescence^31^. In all four PARPi-resistant cell lines, the combination resulted in a statistically significant 2.2-3.4-fold increase in proportion of cells positive for p21, compared with control, with p21 induction significantly greater than that observed with olaparib alone (Figure S2A-D). In contrast, the combination treatment did not increase p21 expression in the parental cell lines. Further, since nuclear area increases in senescence^32^, we compared nuclear sizes across cell lines and treatment conditions and identified up to 16% increased nuclear area due to the combination (Figure S2E,F).

Together, these findings highlight distinct cellular responses in the TNBC cell lines with acquired resistance to PARPi. HCC1395-Res cells, with lower basal nucleolar stress, showed marked NPM1 redistribution following combination of BMH-21 and olaparib, whereas MX1-Res cells demonstrated higher baseline nucleolar stress, and with the combination, demonstrated nucleolar remodeling and less pronounced NPM1 redistribution. All PARPi-resistant cells demonstrated increased DNA damage and p21 expression with the combination treatment, suggesting that the observed synergy in PARPi-resistant cells is associated with nucleolar stress, DNA damage, and senescence-like response.

### Targeting ribosome biogenesis increases PARPi sensitivity in BRCA wild-type TNBC cells

We next extended our analysis to three BRCA wild-type TNBC cell lines with intrinsic PARPi resistance: BT549, Hs578T, and MDAMB231 (Figure 4A-C). We observed synergy between olaparib and BMH-21 in all three cell lines, with CI values ranging from 0.70-0.83. Based on these findings, we then investigated cellular responses to the combination in MDAMB231 cells at 6 and 10 days post-treatment.

**Figure 4.**
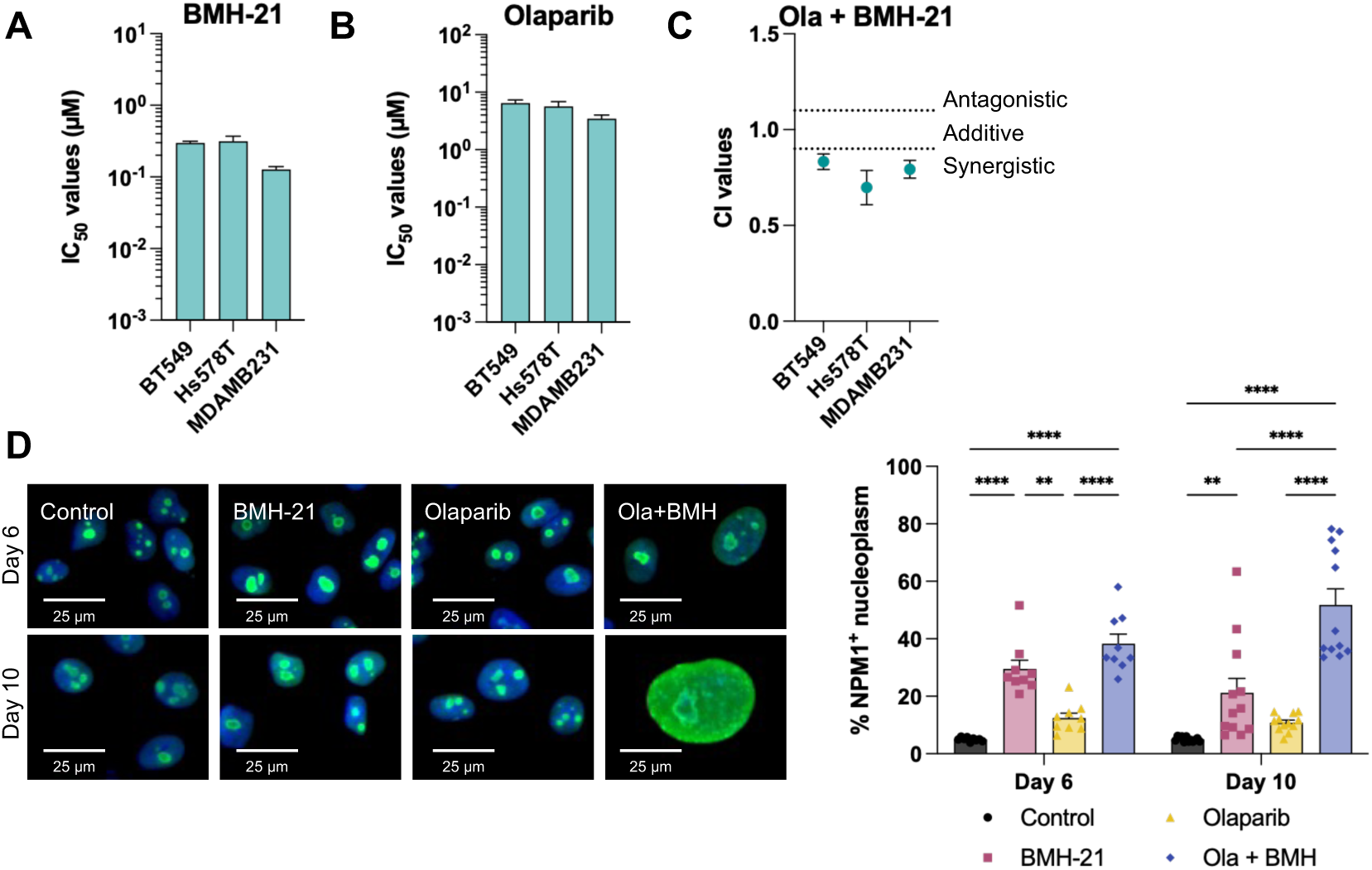
BMH-21 is synergistic with olaparib in cell lines with intrinsic PARPi resistance. IC50 values for **(A)** BMH-21 and **(B)** olaparib in BRCA wild-type TNBC cell lines. **(C)** Combination index values for olaparib and BMH-21 reported at an FA of 0.5. **(D)** Representative images and quantification of NPM1 delocalization in MDAMB231 cells at days 6 and 10 following treatment with DMSO control (black circles), BMH-21 (pink squares), olaparib (yellow triangles), or olaparib combined with BMH-21 (blue diamonds). Images include NPM1 (green) with nuclear DAPI counterstaining (blue). Scale bars indicate 25 µm. Values indicate mean ± SEM. Two-way ANOVA followed by Tukey’s multiple comparisons was performed to test statistical significance within each timepoint, with P < 0.0001****, P <0.001***, P < 0.01**, P < 0.05*.

We first assessed changes in nucleolar morphology in response to single-agent and combination treatments, again using RSL1D1 to identify nucleoli. We identified a 37% increase in nucleolar area on day 10, from 9.4 µm^2^ in control to 12.9 µm^2^ in combination-treated cells, with an associated 31% increase in the number of nucleoli per cell (Figure S3A-C). Interestingly, the increase in nucleolar size was primarily associated with BMH-21, while the number of nucleoli per cell was more dependent on olaparib.

Next, we again evaluated NPM1 localization on days 6 and 10 post-treatment. Similar to HCC1395 resistant cells, we found that the proportion of MDAMB231 cells positive for NPM1 in the nucleoplasm increased by 7.6-and 10.3-fold on days 6 and day 10, respectively, following treatment with the combination of olaparib and BMH-21 (Figure 4D). Together, these data indicate a process of progressive nucleolar remodelling in MDAMB231 cells in response to combined PARPi and ribosome biogenesis inhibition.

### Combined PARPi and ribosome biogenesis inhibition induces a senescence-like phenotype in BRCA wild-type TNBC cells

We then assessed the impact of drug treatment on DNA damage by evaluating γH2AX foci formation in MDAMB231 cells. In comparison to control, the proportion of cells with γH2AX foci formation increased 4.7-and 7.3-fold with the combination of BMH-21 and olaparib on days 6 and 10, respectively (Figure 5A). Next, we used the senescence-associated β-galactosidase (SA-β-gal) assay to determine whether treatment induced a senescence-like phenotype (Figure 5B-D). At day 6, the combination resulted in a 5.6-fold increase in the proportion of SA-β-gal positive cells relative to control, which is concordant with 3.6-fold increase in p21-positive cells observed 6 and 10 days post-treatment with olaparib plus BMH-21. Combination treatment also resulted in significant increases in nuclear area. On day 6 post-treatment, nuclear area increased by 29.0%, from 134.6 µm^2^ in control cells to 173.6 µm^2^ in cells treated with the combination of olaparib and BMH-21. By day 10 post-treatment, nuclear area increased by 52.1%, from 128.8 µm^2^ in control to 195.9 µm^2^ in cells treated with the combination. Overall, the combination had a statistically significant greater impact in comparison to olaparib at 10 days for DNA damage, p21, and nuclear area, and on SA-β-gal positivity at 6 days. Thus, the combination of BMH-21 and olaparib demonstrated increase DNA damage in MDAMB231 cells and are associated with multiple features of a senescence-like response.

**Figure 5.**
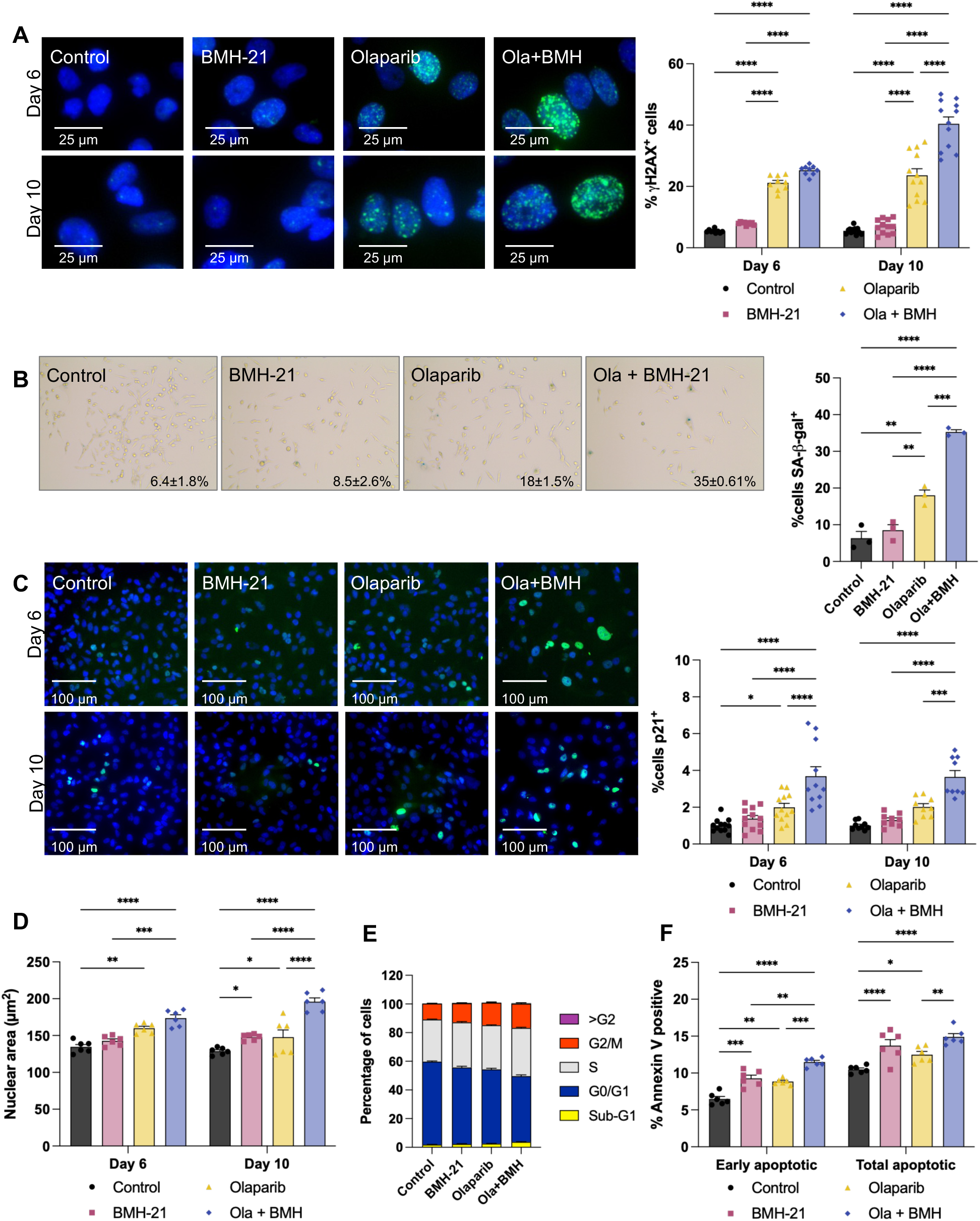
Combined olaparib and BMH-21 induce senescence in TNBC. Representative images and quantification of **(A)** γH2AX, **(B)** SA-β -galactosidase, **(C)** p21, or **(D)** nuclear size in MDAMB231 at day 6 or day 10 post-treatment with DMSO control (black circles), BMH-21 (pink squares), olaparib (yellow triangles), or olaparib combined with BMH-21 (blue diamonds). **(E)** Cell cycle distribution and **(F)** quantification of apoptosis in MDMAB231 cells at day 6 post-treatment. Scale bars indicate **(A)** 25 µm and **(C)** 200 µm. Values indicate mean ± SEM. Two-way ANOVA followed by Tukey’s multiple comparisons was performed to test statistical significance within each timepoint, with P < 0.0001****, P <0.001***, P < 0.01**, P < 0.05*.

Given the concordance of these various senescence markers, we next investigated whether these changes were accompanied by alterations in cell cycle or apoptosis at 6-days post-treatment (Figure 5E, Figure S4). We identified significant increases in the proportions of S-phase cells from 29.4% to 33.7% and G2/M-phase cells from 11.1% to 16.9%, in control and combination-treated cells respectively (Figure 5E, Figure S4B,C). The sub-G1 population also significantly increased to 3.8% in cells treated with olaparib plus BMH-21, relative to 1.9% of control cells. Consistent with the increase in sub-G1 cells, Annexin V staining identified increased apoptosis following combination olaparib and BMH-21 treatment. The proportion of early apoptotic cells increased 1.8-fold from 6.5% of control cells to 11.5% in combination-treated cells, with statistically significant differences with either single agent treatment (Figure 5F). However, total apoptosis was similar between combination cells and BMH-21 alone, suggesting that there may be a temporal difference in the apoptotic responses of cells to these treatments. Together, these findings demonstrate that combined olaparib and BMH-21 treatment in MDAMB231 cells induces a senescence-like response, which is accompanied by altered cell cycle distribution and increased early apoptosis.

### CX-5461 and PARPi combination enhances nucleolar stress and DNA damage in BRCA wild-type TNBC

Another pharmacological approach to target ribosome biogenesis is the Pol I inhibitor CX-5461. Based on the above observations with BMH-21 and previous studies demonstrating synergy with PARPi in pancreatic and ovarian cancers^33,34^, we investigated whether CX-5461 would also synergize with PARPi in TNBC. We used the BRCA wild-type cell lines BT549, Hs578T, and MDAMB231 to measure sensitivity to CX-5461 in a 10-day assay and identified synergy between CX-5461 and olaparib (Figure S5A,B). In agreement with this, we observed a 13.0-fold increase in the proportion of cells with NPM1 redistribution to the nucleoplasm in combination-treated MDAMB231 cells (Figure S5C,D). We identified an 11.2-fold increase in the proportion of γH2AX-positive cells relative to control, and 2.4-4.8-fold increase with either drug alone, along with 3.4-fold increase in cells positive for p21 (Figure S5C,E,F).

Finally, we investigated whether this combination could induce a delayed growth recovery phenotype, as is characteristic of senescence. Following the treatment course, we replated MDAMB231-NuclightRed cells at log-phase density and followed proliferation over three weeks. Control cells reached confluence within a week, and single-agents within two weeks, while combination-treated cells showed sustained proliferative arrest, with relative proliferation remaining essentially unchanged through day 21 (Figure S5G). Together, these data indicate that a second ribosome biogenesis inhibitor, CX-5461, can also induce sensitivity to PARPi via nucleolar stress, DNA damage, and a senescence-like response in BRCA wild-type TNBC cells.

### Ribosome-associated activity correlates with proliferation and DNA repair in treatment-resistant TNBC patients

Having established that inhibition of ribosome biogenesis enhances PARPi sensitivity in TNBC cell lines, we wanted to better understand a clinical context for which the combination of PARPi and ribosome biogenesis could be useful. As part of the Terry Fox Marathon of Hope Cancer Centres Network (MOHCCN)^35,36^, we collected tumour samples from a cohort of TNBC patients who were resistant to neoadjuvant therapy, defined by the presence of a residual tumour at the time of surgery. RNA sequencing of the residual tumours was performed, followed by Gene Set Variation Analysis (GSVA)^37^ to evaluate pathway-level activity between patients (Figure 6A). Given the crosstalk between DNA repair and ribosome biogenesis, along with our previous identification of ribosomal pathways upregulation in PARPi-resistant TNBC, we investigated associations between these pathways in treatment-resistant TNBC. Unsupervised hierarchical clustering showed that patients could be divided into 3 main clusters with pathway enrichment based on 1) core ribosomal and ribosome biogenesis pathways; 2) cell proliferation and senescence pathways; and 3) robust cell proliferation, with upregulation of ribosome biogenesis, DNA repair, and core structural ribosomal pathways in a subset of patients.

**Figure 6.**
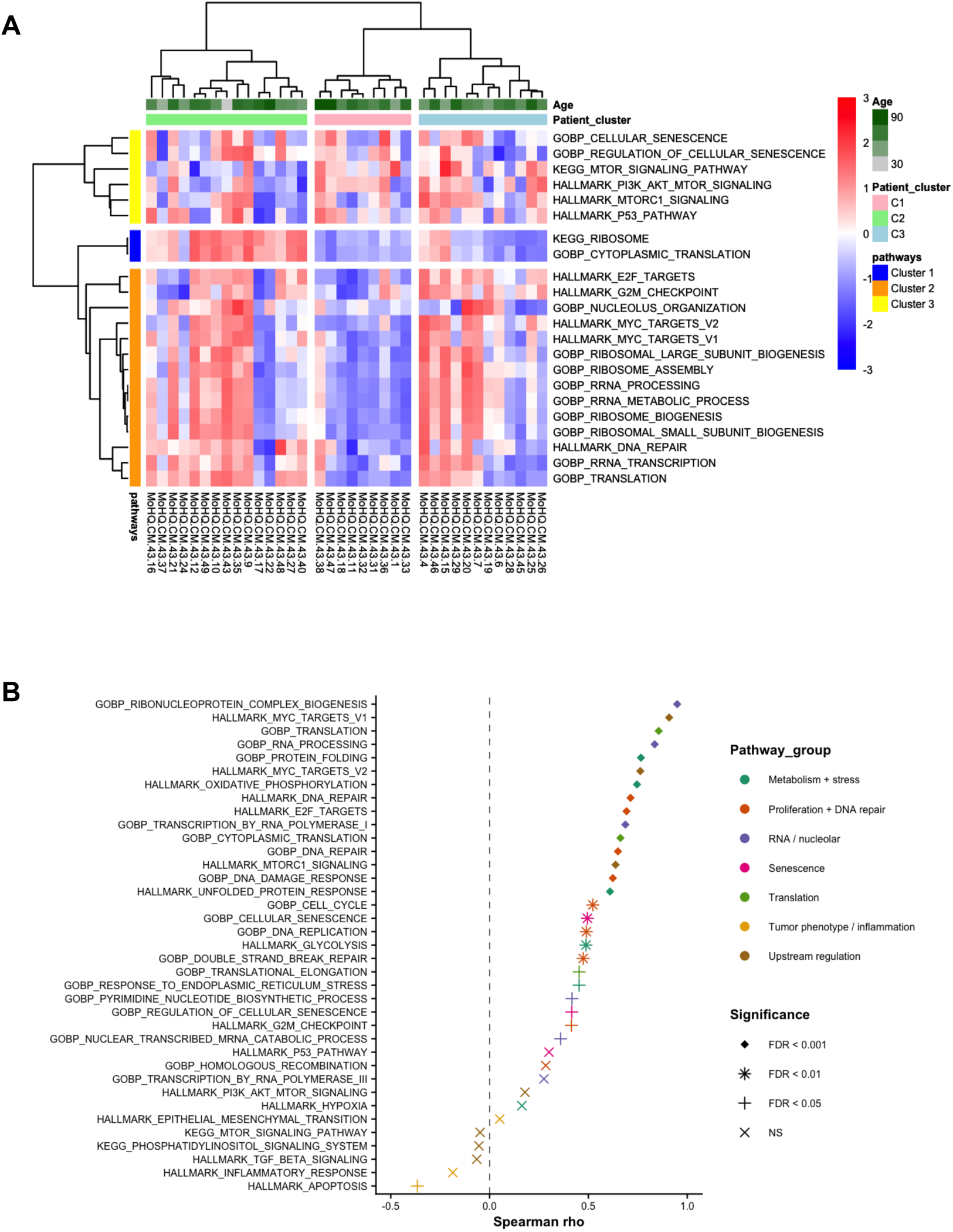
Ribosome pathway activity correlates with proliferation pathways in treatment-resistant TNBC. **(A)** Heatmap with hierarchical clustering of GSVA scores for selected pathways across patients with residual tumors following neoadjuvant treatment. **(B)** Composite ribosome score correlations with biological pathways of interest. Spearman correlations were calculated between mean GSVA score across nine ribosome-associated pathways and selected biological pathways. P-values were adjusted for multiple hypothesis testing by Benjamini-Hochberg procedure, with shape indicating FDR significance. Not significant (NS), X; FDR < 0.05, +; FDR < 0.01, *, FDR < 0.001, diamond.

To examine the cellular processes associated with ribosome activity, we selected gene sets representing ribosome biogenesis, translation, proliferation, DNA damage and repair, and cellular stress pathways (Table S1, Figure S6). We identified significant positive correlations between MYC target gene activity and ribosome biogenesis pathways. In particular, HALLMARK_MYC_TARGETS_V1 and HALLMARK_MYC_TARGETS_V2 positively correlated with seven to nine ribosome-associated pathways (Spearman ρ 0.49-0.92, FDR<0.01). Additionally, several translation-related and proliferative pathways, including GOBP_TRANSLATION, HALLMARK_E2F_TARGETS, and HALLMARK_MTORC1_SIGNALING, were significantly positively correlated with at least seven of the nine ribosome pathways.

DNA repair pathways also showed positive associations with ribosome-associated activity. HALLMARK_DNA_REPAIR and GOBP_DNA_REPAIR were significantly correlated with seven of nine pathways (ρ 0.63-0.76, FDR<0.001), similar to GOBP_DOUBLE_STRAND_BREAK_REPAIR (ρ 0.44-0.60, FDR<0.05, 7/9 pathways). Ribosomal pathways also correlated with DNA damage response and senescence, including GOBP_DNA_DAMAGE_RESPONSE (ρ 0.62-0.76 and FDR<0.001, 7/9 pathways), GOBP_CELLULAR_SENESCENCE (ρ 0.45-0.64, FDR<0.05, 7/9 pathways), and GOBP_REGULATION_OF_CELLULAR_SENESCENCE (ρ 0.37-0.55 FDR<0.05, 3/7 pathways).

Given the substantial overlap between similar and related gene sets, we next tested whether these associations were maintained using a composite ribosome score derived from the same nine ribosome-associated gene sets. The ribosome score identified significant positive correlations with MYC target activity, translation, proliferation, senescence, DNA damage, and DNA repair mechanisms across treatment-resistant samples (Figure 6B). Together, these data indicate a coordinated transcriptional state characterized by high ribosome-associated activity, MYC signalling, translation, proliferation, and DNA repair, supporting ribosome biogenesis as a potential therapeutic vulnerability in early TNBC patients with residual disease.

## Discussion

We are the first to provide a comprehensive overview of the efficacy of combined PARPi and ribosome biogenesis inhibition in TNBC, focusing on a panel of cell lines with acquired or intrinsic resistance to PARPi. Using BRCA-mutant TNBC cell lines with acquired resistance to olaparib and talazoparib and BRCA wild-type TNBC cell lines, we showed synergistic effects with olaparib in combination with BMH-21. Combination treatment was associated with nucleolar stress, DNA damage, and features of senescence-like response. Importantly, these findings were not limited to BMH-21, as combining olaparib with the ribosome biogenesis inhibitor CX-5461 produced similar synergistic effects in MDAMB231 cells. Finally, we identified correlations between ribosome-associated pathways and proliferation programs in a cohort of treatment-resistant TNBC patients. Together, these data support the combination of PARPi and ribsosome biogenesis inhibition as a targeted therapeutic combination strategy in PARPi-resistant breast cancer and the clinical significance of ribosome biogenesis in TNBC patients.

Although PARPi were developed based on the role of PARP-1 in the DNA damage response, PARP-1 also has important functions in the nucleolus, where it contributes to rDNA transcription and ribosome biogenesis^15,20^. In the nucleolus, PARP-1 interacts with small nucleolar (sno)RNAs and promotes PARylation of the DEAD-box RNA helicase 21 (DDX21), which supports rDNA transcription and proliferation in breast cancer cells^38,39^. Consistent with this connection between PARP-1 and ribsosome biogenesis, PARPi-induced redistribution of DDX21 and reductions in rRNA synthesis have been reported in endometrial and ovarian cancer^40^. These findings provide a potential mechanistic link between PARPi and ribosome biogenesis and support our observation that PARPi-resistant cells demonstrate an altered nucleolar organization.

We demonstrated a transcriptional upregulation of ribosome pathways in 3 of 4 cell lines with acquired resistance to PARPi. Our findings are consistent with previous studies implicating ribosome-related pathways in PARPi resistance. In a BRCA2-deficient mouse model with acquired PARPi resistance, multi-omic analysis identified KEGG_Ribosome as the most highly upregulated pathway, while an unbiased CRISPR screen in a PARG-deficient, BRCA-deficient model identified ribosome biogenesis, RNA degradation, and RNA transport among resistance-associated pathways^41,42^. Ribosomal protein S6 phosphorylation has also been implicated in resistance to BRCA1-deficient tumors^43^. The variable detection of these pathways across previous screens may reflect the heterogeneity and context dependence of ribosome biogenesis, as well as differences in screening approaches. In particular, complete loss of essential ribosome biogenesis factors may be poorly tolerated, limiting their identification in knockout screens, while short-term screens may miss vulnerabilities that manifest through sustained proliferative arrest^44,45^. Our findings support this context dependence, with transcriptional activation of ribosome-related pathways observed in a subset of our PARPi-resistant cell lines, and in a subset of treatment-resistant TNBC patient tumours.

Our results highlight that TNBC cell lines with acquired resistance to PARPi can have two distinct baseline phenotypes yet still achieve synergy in combination with Pol I inhibition. PARPi-resistant HCC1395 cells demonstrated reductions in nucleolar area and fewer nucleoli per cell, together with reduced NPM1 redistribution, consistent with lower baseline nucleolar stress. Following combination treatment, these cell lines demonstrated marked NPM1 redistribution, indicating induction of nucleolar stress. In contrast, PARPi-resistant MX1 cells demonstrate higher baseline nucleolar stress, evidenced by larger nucleolar area, increased number of nucleoli, and greater NPM1 redistribution. With the combination treatment, these cell lines demonstrated nucleolar remodeling and NPM1 relocalization was less pronounced than those observed in HCC1395-resistant cells. Despite these differences in baseline and treatment-induced nucleolar stress, both models remained sensitive to the combination of olaparib and Pol I inhibition. Interestingly, nucleolar stress has recently been characterized using a gene signature that stratified tumorus into functional states and was associated with clinical outcomes^46^.

Combination treatments with either BMH-21 or CX-5461 and olaparib was synergistic in BRCA-mutant PARPi-resistant cell lines, and three BRCA wild-type TNBC cell lines, with increased nucleolar stress, sustained DNA damage, and senescence-like phenotypes in MDAMB231 cells. CX-5461 has previously been investigated in combination with PARPi in ovarian^34^ and prostate^47^ cancers, where it was also shown to increase DNA damage in combination with PARPi in the contexts of acquired and intrinsic PARPi resistance, respectively. Given the established links between persistent DNA damage, nucleolar stress, and senescence^15,18,19,48,49^, our findings indicate that disruption of ribosome biogenesis may contribute to the induction of a senescence-like state following PARPi treatment. Further, comparison of the 6-and 10-day timepoints suggests the accumulation of these stresses over the course of treatment. Notably, MDAMB231 cells lack functional p53, indicating that the combination of ribosome biogenesis inhibition with PARPi may induce senescence independent of canonical p53-mediated signaling, consistent with previous studies^18^.

The best approach to target ribosome biogenesis is still under investigation. CX-5461 is currently in 5 clinical trials for various cancers, including Phase I studies in patients with advanced solid tumours enriched for DNA-repair deficiencies (NCT04890613) and in combination therapy for HER2-positive breast cancer (NCT07137416). Despite being well studied, CX-5461 also has activities beyond ribosome biogenesis, inducing replicative stress, DNA damage, and acts as a G-quadruplex stabilizer in cellular models^33,34,50^. We therefore focused on BMH-21, as it does not induce DNA damage.

To better understand the potential clinical utility of combining ribosome biogenesis inhibition and PARPi, we evaluated a cohort of treatment-resistant TNBCs. Hierarchical clustering divided the cohort into three main groups, with patients demonstrating increased activity of ribosomal and ribosome biogenesis pathways, together with proliferative and senescence-associated programs. These findings were similar to the transcriptional heterogeneity observed across our PARPi-resistant cell lines. Together, these findings demonstrate that increased ribosomal pathway activity is present in a subset of treatment-resistant TNBCs, and support further investigation for ribosome biogenesis inhibition as a therapeutic strategy in this setting.

Taken together, our results identify ribosome biogenesis as a potential therapeutic vulnerability in PARPi-resistant TNBC. Cell lines with acquired resistance to PARPi demonstrate two distinct baseline nucleolar states categorized by lower and higher levels of nucleolar stress yet four such cell lines demonstrated synergy with PARPi and Pol I inhibition. We also demonstrated efficacy of the combination in BRCA wild-type TNBC cell lines with intrinsic resistance to PARPi accompanied with DNA damage, senescence, and cell death. Together, these findings support further investigation of ribosome biogenesis inhibitors in combination with PARPi in PARPi-resistant and treatment-resistant TNBC.

## Methods

### Cell culture

MDAMB231 and MDAMB231-NuclightRed breast cancer cells were cultured in DMEM (Gibco, 11995-073) supplemented with 10% FBS (Gibco, A5670701). HCC1395 and MX1 parental and resistant cell lines were cultured in RPMI (Gibco, 11875-119) supplemented with 10% FBS. All cells were grown at 37°C with 5% CO_2_.

PARPi-resistant MX1 cells were derived from MX1 TNBC cells grown in RPMI with increasing concentrations of talazoparib or olaparib added to the media over the course of six months, as previously described^51^. Cells were passaged in 6-well tissue culture-treated plates and allowed to adhere. The next day, media containing 0.0006 μM talazoparib or 0.14 μM olaparib was added. Cells were treated for rounds of 3-5 days, then allowed to recover to near confluence in complete media lacking PARPi. Media was exchanged every 3 to 5 days, alternating between media containing or lacking PARPi depending on cell growth. Treatment with PARPi was repeated at the same concentration until resistance was observed, at which point PARPi concentration was doubled. This process was repeated until sufficient resistance was observed. Finally, stable resistance was confirmed by IC50 calculation after 5 passages in the absence of PARPi and a freeze-thaw cycle.

### RNA extraction and sequencing of cell lines

MX1-Parental, -OlaR, and -TalaR cells were seeded at 50 000 cells per mL in 6-well tissue culture plates and grown for four days. Cells were harvested in 500 µL of Trizol (Invitrogen, 15596018) per well, combining two wells per condition and 3 independent samples, then stored at −80°C until RNA extraction. RNA was extracted using RNeasy Plus mini kit (Cat. #74134), with a modified phenol-chloroform extraction prior to sample loading on the columns. RNA was quantified using Qubit (Thermo Scientific) and quality was assessed with the 2100 Bioanalyzer (Agilent Technologies). Libraries were prepared and samples were sequenced with Illumina NextSeq 2000, using the Illumina kit NextSeq 75 cycles High Output v2, with ∼ 50 M reads per sample, at the Institute for Research in Immunology and Cancer (IRIC, Montreal).

The first part of the analysis including normalization and DeSEq2 analysis was performed at the Bioinformatics Facility at IRIC. Sequences were trimmed for sequencing adapters and low quality 3’ bases using Trimmomatic version 0.38^52^ and aligned to the reference human genome version GRCh38 (gene annotation from Gencode version 40, based on Ensembl 106) using STAR version 2.7.1a^53^. Gene expressions were obtained as readcount directly from STAR as well as computed using RSEM^54^ in order to obtain normalized gene and transcript level expression, in TPM values, for these stranded RNA libraries. DESeq2 version 1.30.1^55^ was then used to normalize STAR gene readcounts and produce sample clustering. Using the log2-fold change values from the DESeq2 analysis, all expressed genes were ranked and a pre-ranked Gene Set Enrichment Analysis was performed using GenePattern^56^. Gene sets from MsigDB were used, including Hallmarks and KEGG legacy for human tissue. FDR cutoff of 25 % and p-value < 0.05 were used to identify statistically significant gene sets. Dot plots were created in RStudio (version 2026.08.1+195).

### IC50 and Combination Index calculations

IC50 and CI values were calculated in a 10-day assay, as previously^14^, using the Chou and Talalay approach^28^. Briefly, cells were seeded on day 0, then treated with 3-fold dilutions (0.0015 to 10 µM) of BMH-21 (SelleckChem, S7718-10MG) or 5-fold dilutions (0.00026 to 100 µM) of olaparib (SelleckChem, S1060-100MG) on day 1. On day 4-5 cell media and drugs were refreshed, and on day 10 cells were fixed for 15 minutes in 4% paraformaldehyde (PFA), washed in PBS, and stained with DAPI for 30 minutes to 1 hour. Cells were washed 3 times with PBS and imaged at 20X on Operetta CLS (Revvity), with 37 images per well. Cells were identified and counted with Harmony High-Content Imaging and Analysis Software (Version 5.2, Revvity).

For CI calculations, 2-fold dilutions of BMH-21, olaparib, or the combination were centred around the IC50 for each cell line. Treatments and media changes were timed the same as above, then followed with cell fixation, DAPI staining, and imaging as described. CI values were calculated using CompuSyn (ComboSyn Incorporated, Paramus, NJ, USA) at Fa = 0.5.

### Immunofluorescence and high-content imaging

Cells were seeded on 96-well plates on day 0. The next day, cells were treated with drugs at 10-day IC_50_ concentrations. Cells were grown until fixation on day 6 or day 10, with a media and drug refresh on day 4-5. Cells were fixed for 15 minutes in 4% PFA, then washed three times in PBS and stored in 4 °C or permeabilized immediately in 0.3% Triton X-100 and 2% bovine serum albumin (BSA) in PBS for 45 minutes at room temperature. Incubation with primary antibodies diluted in 2% BSA in PBST followed overnight at 4 °C. Finally, cells were washed three times with PBST, incubated with secondary antibodies for 1-2 hours at room temperature, and washed a final two times each with PBST and PBS. Plates were imaged immediately or stored in 4 °C until imaging. All primary and secondary antibodies can be found in Supplementary Table S2.

Fix and stained cells were imaged on the Operetta CLS (PerkinElmer) with 20X or 40X objectives. Filters were specific for Alex 488, Alexa 647, or DAPI and 37 or 105 images were scanned per well at 20X and 40X respectively. Image analysis was performed using Harmony High-Content Imaging and Analysis Software (Version 5.2, PerkinElmer).

For single cell analysis, nucleoplasmic NPM1 intensity in parental or DMSO-treated control cells was used to establish a cutoff corresponding to 5% of cells exhibiting NPM1 delocalization. Similarly, as previously^14^, the number of γH2AX foci in control cells was used to establish a 5% cutoff for γH2AX positivity, while a 1% cutoff was used for identifying p21 positivity. These analyses were followed by comparison of the proportions of NPM1, γH2AX, or p21 positive cells across treatment conditions.

### Senescence-associated β-galactosidase staining

SA-β-Gal activity was assessed using an adapted protocol^57^. Cells were seeded in 6-well plates on day 0 and treated on day 1. On day 6, cells were washed twice with PBS and fixed with 0.5% glutaraldehyde in PBS for 15 minutes at room temperature. Cells were washed with PBS and incubated at 37 °C in the absence of CO₂ with freshly prepared SA-β-Gal staining solution (1 mg/mL X-Gal, 40 mM citric acid/sodium phosphate buffer (pH 6.0), 150 mM NaCl, 2 mM MgCl_2_, 5 mM potassium ferricyanide, and 5 mM potassium ferrocyanide). Staining solution was filtered through a 0.2-µm filter before use and 1.5 mL used per well in 6-well plates. Cells were incubated for 2 to 6 hours protected from light, with staining monitored periodically to avoid excess background. Reactions were stopped upon visible blue staining in a minimal proportion of control cells. Cells were washed twice with PBS and maintained in PBS until imaging. For quantification, SA-β-Gal-positive and total cells were manually counted from representative microscopic fields using the Cell Counter plugin in Fiji (ImageJ), with a minimum of 200 cells counted per condition.

### Cell cycle and apoptosis analysis by flow cytometry

MDAMB231 cells were seeded on 6-well plates on day 0 and treated on day 1. After incubation under the indicated experimental conditions, cells were collected on day 6, washed with ice-cold PBS, and fixed/permeabilized in 70% ethanol at −20 °C overnight. Subsequently, cells were washed with ice-cold PBS and stained with 50 µg/mL propidium iodide (PI) DNA staining solution containing 200 µg/ml RNase A (ThermoFisher, R1253) and 0.2% (v/v) Triton X-100 in PBS. Cells were incubated for 1 hour at room temperature in the dark and analyzed by flow cytometry using a BD FACSymphony A3 flow cytometer (BD Biosciences). A minimum of 10 000 cells were acquired for each sample and cell cycle phase distribution was determined using FlowJo software (version 10.10.1).

Apoptosis was assessed by Annexin V-FITC/PI double staining method. Briefly, on day 6, adherent cells were harvested, washed with cold PBS, and resuspended at a concentration of 1 million cells per mL in Annexin V binding buffer (10 mM HEPES, 140 mM NaCl, and 2.5 mM CaCl_2_, adjusted to pH 7.4). Annexin V-FITC (Invitrogen, A13199) was added to cells at a ratio of 1:200 and PI at a final concentration of 1 µg/ml, followed by incubation in the dark at room temperature for 15 minutes. Samples were analyzed within 30 minutes of staining on a BD FACSymphony A3 flow cytometer (BD Biosciences). Data were analyzed using FlowJo software, where cells negative for both Annexin V and PI were considered viable, Annexin V-positive/PI-negative cells were classified as early apoptotic, and Annexin V-positive/PI-positive cells were classified as late apoptotic.

### Growth recovery assay

MDAMB231-NuclightRed cells were plated in 10 cm plates with 10 mL of full DMEM and treated with IC50 concentrations of CX-5461, olaparib, or the combination, according to the 10-day assay. On day 10, cells were trypsinized, counted, and replated at 25 000 cells/mL in 96-well black plates with three wells per condition. Cells were imaged every second day using Incucyte S3 (Sartorius) with phase and red filters. Analysis was performed in Incucyte 2026A software, using red objects to count cell numbers. Cell numbers were normalized to the first image, 24 hours following drug removal and replating.

### RNA-sequencing and pathway analysis of patient samples

RNA-sequencing data from 36 TNBC patients were included in the Marathon of Hope Québec – *Centre Hospitalier de l’Université de Montréal* – 43 (MoHQ-CM-43) residual disease cohort of the Terry Fox Marathon of Hope Canadian Cancer Network^35,36^. This project and the biobank were approved by our institutional research ethics board (21.137). FFPE-preserved tissue was used to RNA extraction using RNeasy FFPE Kit (Qiagen) in combination with the Qiacube according to protocol. From FFPE blocks, 2 sections were used for groups with >70% cellularity, and 4 sections for groups between 30-70% cellularity. Quality control was performed by TapeStation RNA ScreenTape (Agilent).

Whole-transcriptome sequencing was performed using a minimum of 80M paired end reads. Data were analyzed in R Studio (version 2026.08.1+195) using transcript-level transcripts-per-million (TPM) estimates. Transcript identifiers were mapped to HGNC gene symbols using the org.Hs.eg.db annotation package^58^ and TPM values were summed for transcripts mapping to the same gene. Genes with TPM greater than 1 in at least 20% of samples were retained, followed by transformation using log2(TPM + 1).

Gene set variation analysis (GSVA) was performed using the GSVA package^37^. Gene sets were obtained from the Molecular Signatures Database (MSigDB) using the msigdbr package^59^, including Hallmark, KEGG Legacy, and Gene Ontology Biological Process gene sets^60,61^. Selected gene sets were analyzed by GSVA using a Gaussian kernel with gene-set sizes restricted to between 5 and 500 genes. Selected GSVA scores were visualized for pathway activity across patients using row-scaled heatmaps. Patients and pathways were hierarchically clustered using the correlation distance (1-Pearson correlation) and Ward D2 method, with three clusters specified for each dimension.

To examine relationships between ribosome-associated pathways and other biological processes, Spearman rank correlations were calculated between individual ribosome-associated pathway enrichment scores and all other selected pathway scores. P-values were adjusted for multiple testing using the Benjamini-Hochberg procedure. Finally, a composite ribosome score was calculated as the mean GSVA score across nine predefined ribosome-associated pathways and Spearman correlations between the composite score and individual non-ribosome pathway scores were calculated with P-values adjusted using the Benjamini-Hochberg procedure.

## Acknowledgements

We thank the Cellular Imaging, Cellular Physiology, and Molecular Pathology platforms of the Centre de Recherche de Centre hospitalier de l’Université de Montréal (CRCHUM), especially Véronique Barrès and Liliane Meunier. We thank Raphaelle Lambert and Patrick Gendron from the Genomics and Bioinformatics Facility of the Institut de Recherche en Immunologie et Cancérologie (IRIC) for the RNA-Seq analysis.

We would like to acknowledge Pascale Marquis and her colleagues at McGill University’s Canadian Centre for Computational Genomics (C3G) for their work in performing secondary bioinformatics analyses and quality control of the whole-transcriptome sequencing data used in this study.

## CRediT authorship contribution statement

**Mallory I. Frederick:** Writing – review & editing, Writing – original draft, Visualization, Methodology, Investigation, Formal analysis. **Nicolas Pouderous:** Writing – review & editing, Visualization, Methodology, Investigation, Formal analysis. **Kenan Aloss:** Methodology, Investigation, Formal analysis. **Qi Yang:** Investigation, Formal analysis. **Gerardo Ferbeyre:** Writing – review & editing, Supervision, Funding acquisition, Conceptualization. **Saima Hassan:** Writing – review & editing, Supervision, Project administration, Funding acquisition, Conceptualization.

## Supplementary Figure Legends

**Supplementary Figure S1.**
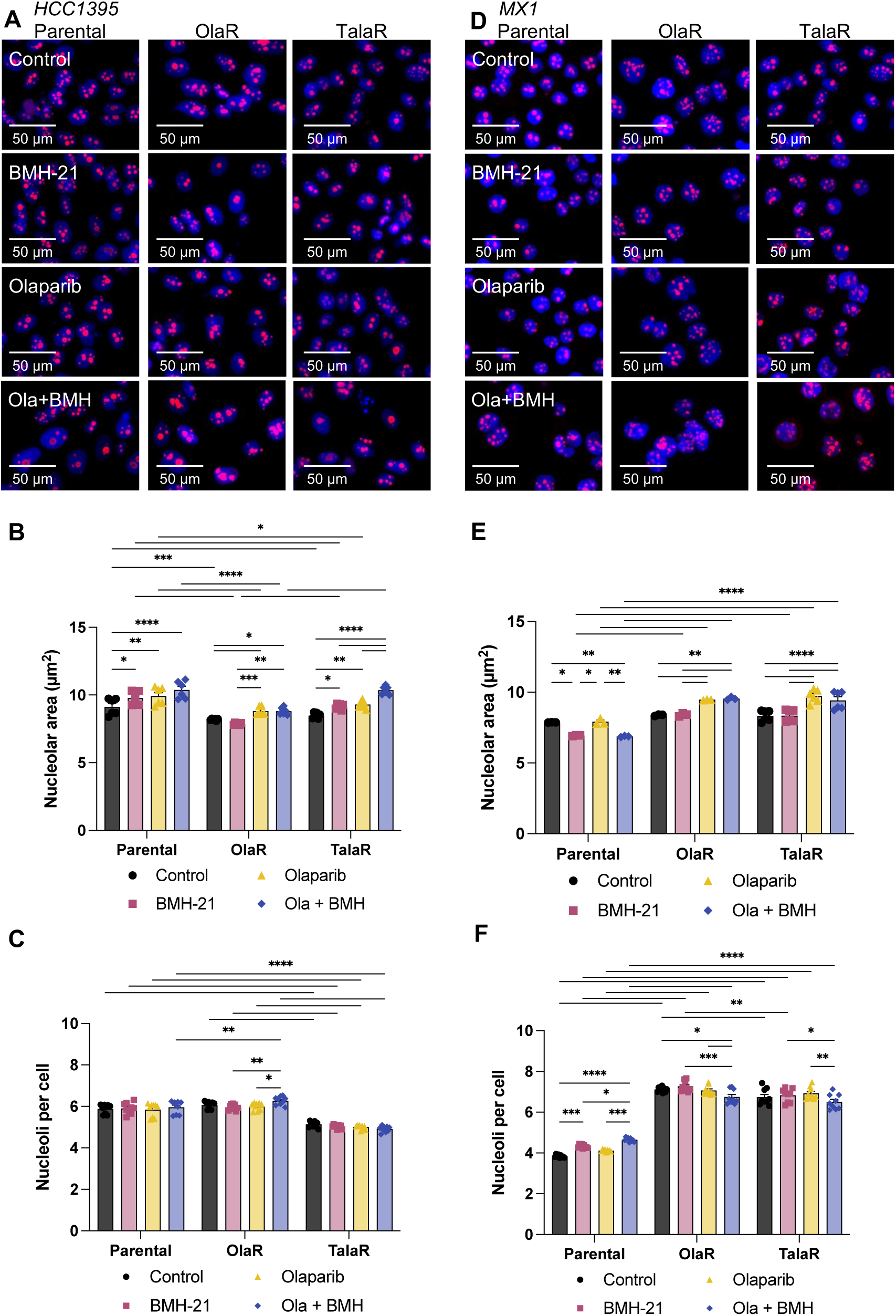
Nucleolar morphology is altered in response to combined BMH-21 and olaparib in acquired PARPi resistance. Representative immunofluorescence images of **(A)** HCC1395 and **(D)** MX1 parental and resistant cell lines treated with DMSO control, BMH-21, olaparib, or olaparib combined with BMH-21. Images include RSL1D1 (red) with nuclear DAPI counterstaining (blue). Scale bars indicate 50 µm. Quantification of nucleolar area in **(B)** HCC1395 and **(E)** MX1 and number of nucleoli in **(C)** HCC1395 and **(F)** MX1 parental and acquired PARPi-resistant cell lines following treatment with DMSO control (black circles), BMH-21 (pink squares), olaparib (yellow triangles), or olaparib combined with BMH-21 (blue diamonds). Bar graphs indicate mean ± SEM. Two-way ANOVA followed by Tukey’s multiple comparisons was performed to test statistical significance within each cell line, with P < 0.0001****, P <0.001***, P < 0.01**, P < 0.05*.

**Supplementary Figure S2.**
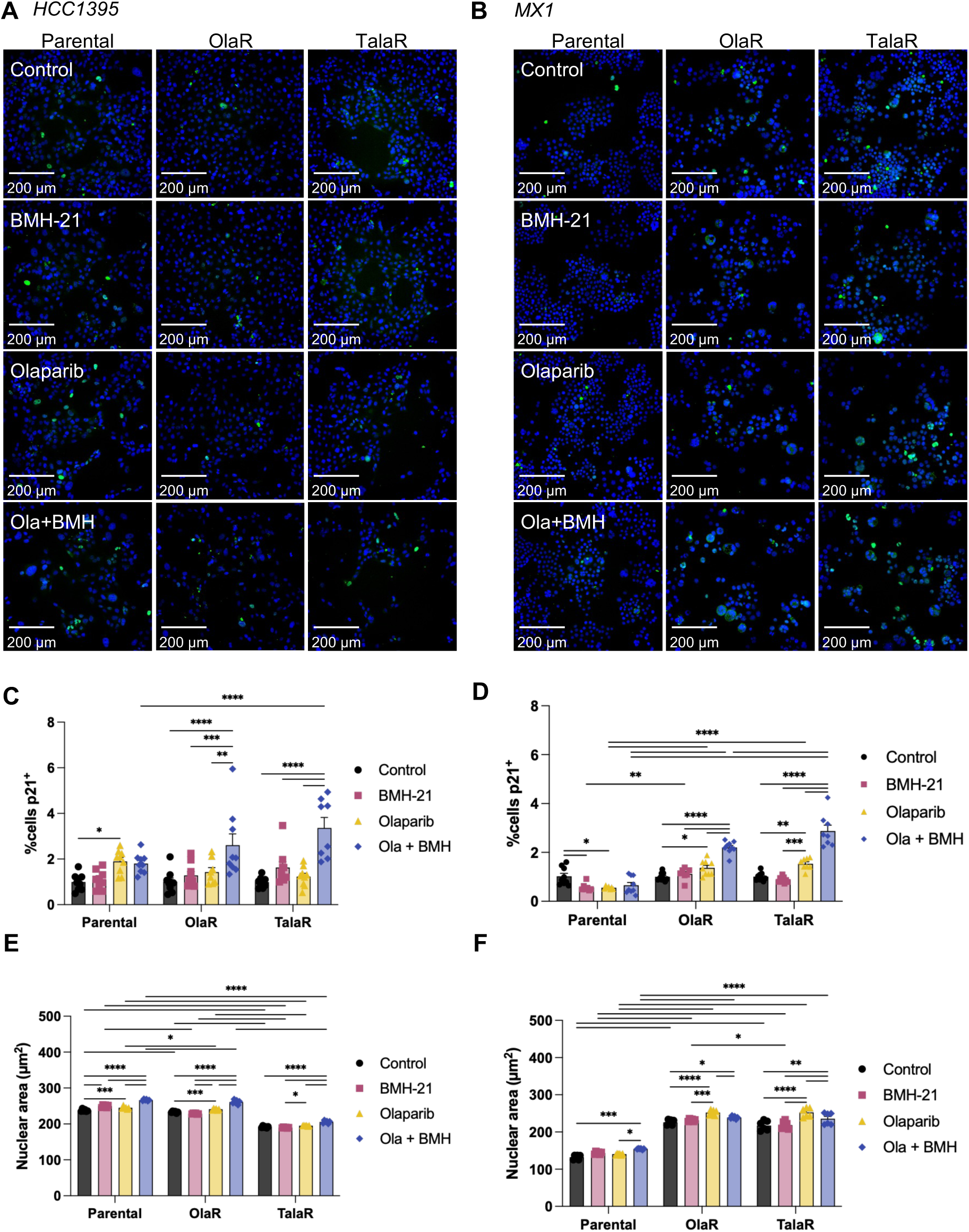
Combined BMH-21 and olaparib induces senescence-associated changes in PARPi-resistant TNBC. **(A,B)** Representative immunofluorescence images and **(C,D)** quantification of p21 positive cells in **(A,C)** HCC1395 and **(B,D)** MX1 parental and resistant cell lines treated with DMSO control (black circles), BMH-21 (pink squares), olaparib (yellow triangles), or olaparib combined with BMH-21 (blue diamonds). Images include p21 (green) with nuclear DAPI counterstaining (blue). Scale bars indicate 200 µm. Quantification of nuclear area in **(E)** HCC1395 and **(F)** MX1 cell lines treated as in (A-D). Bar graphs indicate mean ± SEM. Two-way ANOVA followed by Tukey’s multiple comparisons was performed to test statistical significance within each cell line, with P < 0.0001****, P <0.001***, P < 0.01**, P < 0.05*.

**Supplementary Figure S3.**
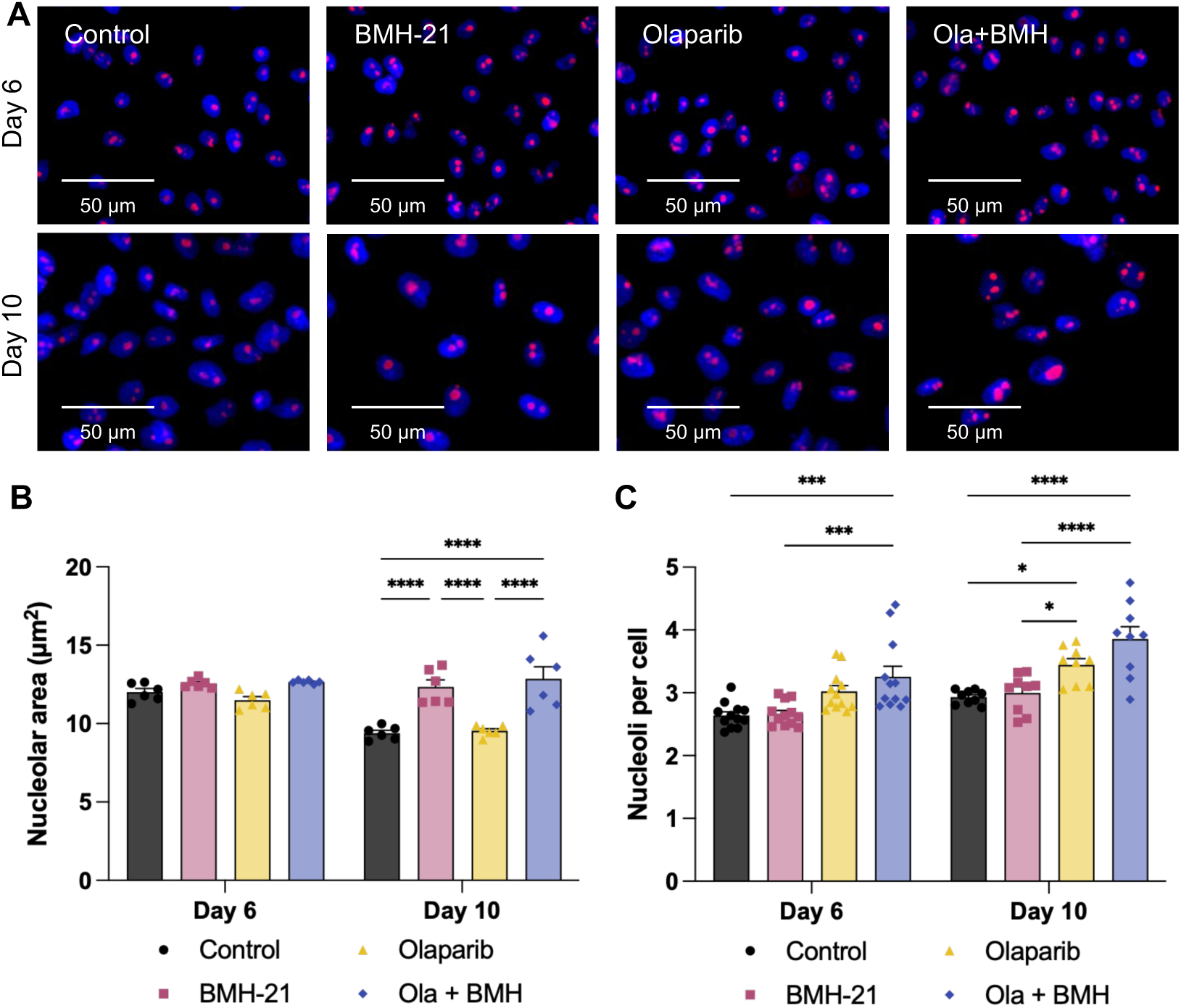
Combined BMH-21 and olaparib induce temporal changes in nucleolar morphology. **(A)** Representative immunofluorescence images of RSL1D1 (red) staining in MDAMB231 cells, with nuclear DAPI counterstaining (blue). Scale bars indicate 50 µm. Quantification of **(B)** nucleolar area and **(C)** number of nucleoli on day 6 or day 10 following treatment with DMSO control (black circles), BMH-21 (pink squares), olaparib (yellow triangles), or olaparib combined with BMH-21 (blue diamonds). Two-way ANOVA followed by Tukey’s multiple comparisons was performed to test statistical significance within each timepoint, with P < 0.0001****, P <0.001***, P < 0.01**, P < 0.05*.

**Supplementary Figure S4.**
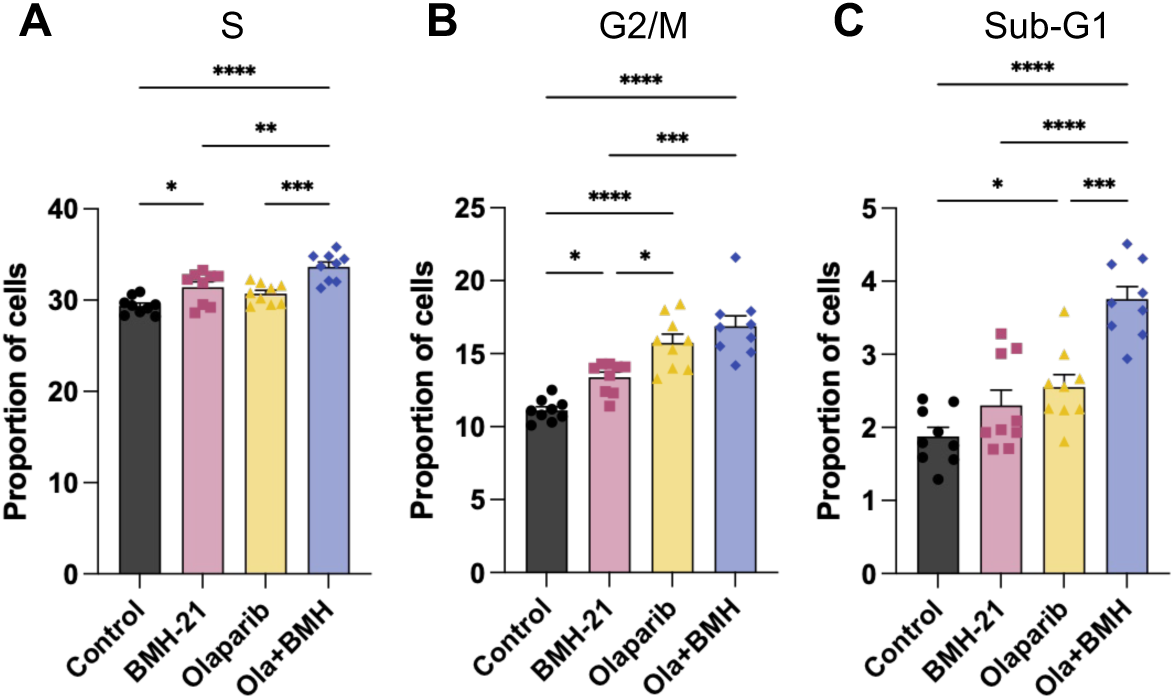
Combined BMH-21 and olaparib alter cell cycle distribution in MDAMB231 cells. Quantification of the percentage of cells in **(A)** S, **(B)** G2/M, and **(C)** sub-G1 phases following treatment with DMSO control (black circles), BMH-21 (pink squares), olaparib (yellow triangles), or olaparib combined with BMH-21 (blue diamonds). Bar graphs indicate mean ± SEM. One-way ANOVA followed by Tukey’s multiple comparisons was performed for each cell cycle phase, with P < 0.05*, P < 0.01**, P < 0.001***, P < 0.0001****.

**Supplementary Figure S5.**
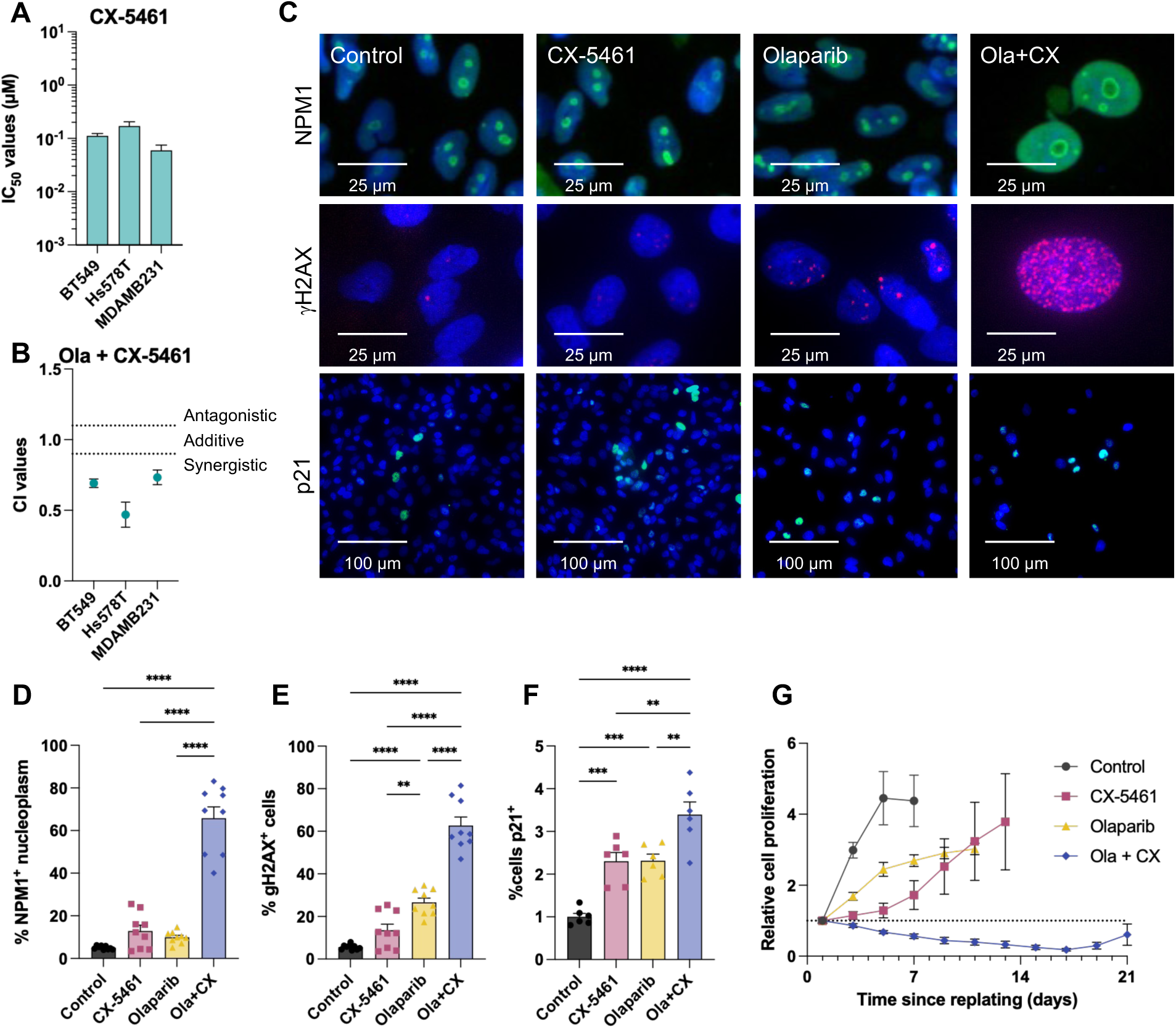
CX-5461 synergizes with olaparib to induce a senescence-like phenotype in MDAMB231 cells. **(A)** IC50 values for CX-5461 in BRCA wild-type TNBC cell lines. **(B)** Combination index values for olaparib and CX-5461 reported at an FA of 0.5. **(C)** Representative immunofluorescence images and **(D-F)** quantification of **(D)** NPM1 delocalization, **(E)** γH2AX foci formation, and **(F)** p21 expression in MDAMB231 cells treated with DMSO control (black circles), CX-5461 (pink squares), olaparib (yellow triangles), or olaparib combined with CX-5461 (blue diamonds). Scale bars indicate 25 µm (NPM1, γH2AX) or 100 µm (p21). **(G)** Quantification of relative cell proliferation until confluence following 10-day treatments as in (C-F). Graphs indicate mean ± SEM. One-way ANOVA followed by Tukey’s multiple comparisons was performed, with P < 0.05*, P < 0.01**, P < 0.001***, P < 0.0001****.

**Supplementary Figure S6.**
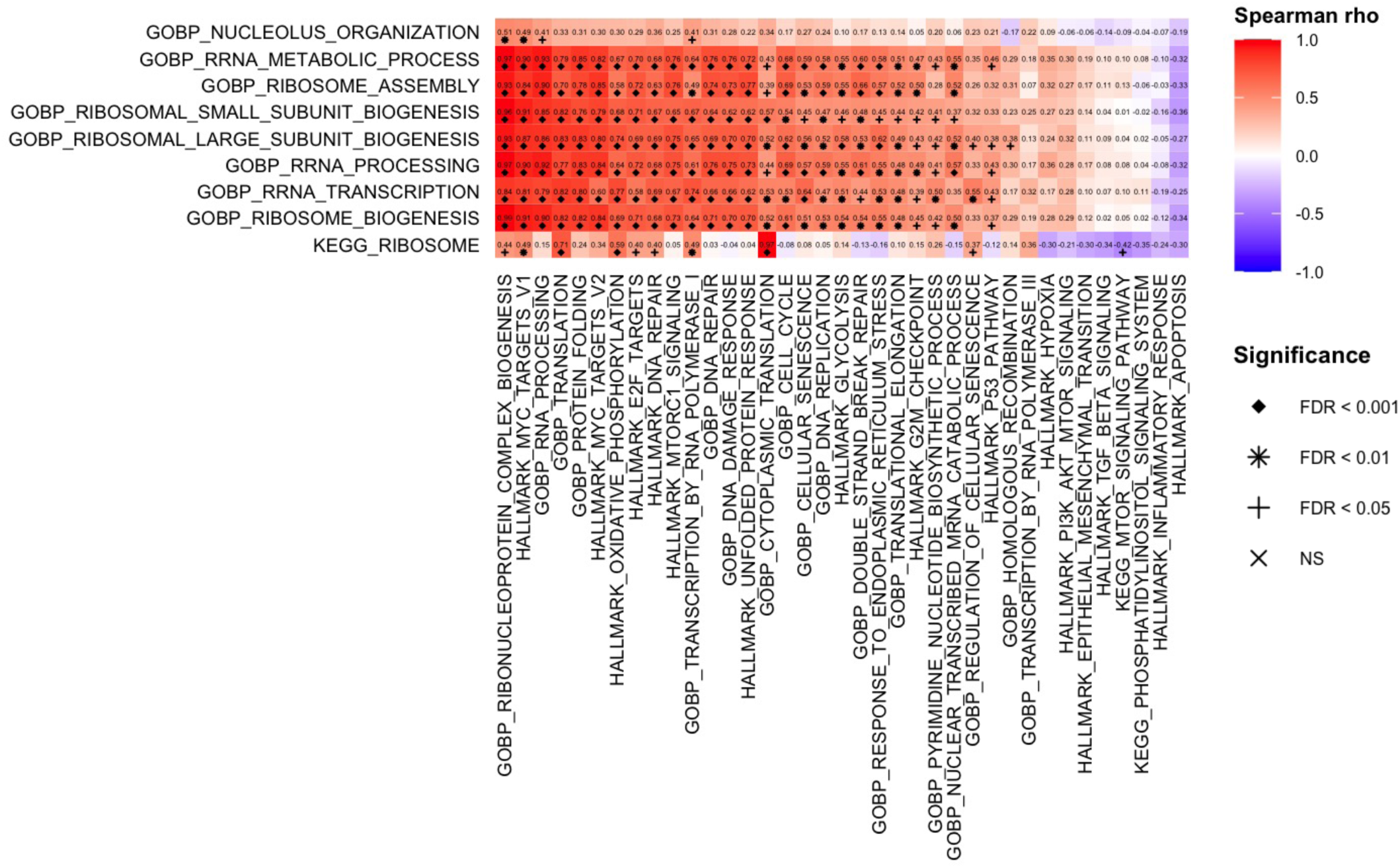
Correlation between individual ribosome-associated pathways and selected biological processes in treatment-resistant TNBC patients. Spearman correlations between GSVA scores for nine ribosome-associated pathways and selected pathways of interest in patients with residual tumors following neoadjuvant treatment. P-values were adjusted for multiple hypothesis testing by Benjamini-Hochberg procedure, with shape indicating FDR significance. FDR < 0.05, +; FDR < 0.01, *, FDR < 0.001, diamond.

