## Supplemental Tables for "Targeting ribosome biogenesis sensitizes PARPi-resistant triple-negative breast cancer"

**Table S1.** Pathways used in ribosome targeted GSEA analysis of HCC1395 and MX1 cell lines.

| **Pathway** | **Number of genes** |
|---|---|
| KEGG_RIBOSOME | 88 |
| GOBP_RIBOSOME_BIOGENESIS | 334 |
| GOBP_RRNA_PROCESSING | 236 |
| GOBP_NUCLEOLAR_LARGE_RRNA_TRANSCRIPTION_BY_RNA_POLYMERASE_I | 33 |
| GOBP_RIBOSOMAL_LARGE_SUBUNIT_BIOGENESIS | 80 |
| GOBP_RIBOSOME_ASSEMBLY | 71 |
| GOBP_NUCLEOLUS_ORGANIZATION | 18 |
| GOBP_TRANSCRIPTION_BY_RNA_POLYMERASE_I | 48 |
| GOBP_TRANSCRIPTION_BY_RNA_POLYMERASE_III | 33 |
| GOBP_REGULATION_OF_TRANSCRIPTION_OF_NUCLEOLAR_LARGE_RRNA_BY_RNA_POLYMERASE_I | 15 |
| GOBP_TRANSCRIPTION_ELONGATION_BY_RNA_POLYMERASE_I | 16 |
| GOBP_TRANSCRIPTION_INITIATION_AT_RNA_POLYMERASE_I_PROMOTER | 28 |
| HALLMARK_MYC_TARGETS_V1 | 200 |
| HALLMARK_E2F_TARGETS | 200 |
| HALLMARK_MTORC1_SIGNALING | 200 |

**Table S2.** Pathways used in targeted GSVA analysis of patient residual tumors.

| ***Ribosome / ribosome biogenesis*** | ***Number of genes*** |
|---|---|
| KEGG_RIBOSOME | 88 |
| GOBP_RIBOSOME_BIOGENESIS | 334 |
| GOBP_RRNA_TRANSCRIPTION | 47 |
| GOBP_RRNA_PROCESSING | 236 |
| GOBP_RIBOSOMAL_LARGE_SUBUNIT_BIOGENESIS | 80 |
| GOBP_RIBOSOMAL_SMALL_SUBUNIT_BIOGENESIS | 114 |
| GOBP_RIBOSOME_ASSEMBLY | 71 |
| GOBP_RRNA_METABOLIC_PROCESS | 289 |
| ***RNA / nucleolar biology*** | |
| GOBP_NUCLEOLUS_ORGANIZATION | 18 |
| GOBP_RNA_PROCESSING | 1484 |
| GOBP_RIBONUCLEOPROTEIN_COMPLEX_BIOGENESIS | 530 |
| GOBP_NUCLEAR_TRANSCRIBED_MRNA_CATABOLIC_PROCESS | 148 |
| GOBP_TRANSCRIPTION_BY_RNA_POLYMERASE_I | 75 |
| GOBP_TRANSCRIPTION_BY_RNA_POLYMERASE_III | 62 |
| ***Translation*** | |
| GOBP_TRANSLATION | 766 |
| GOBP_CYTOPLASMIC_TRANSLATION | 177 |
| GOBP_TRANSLATIONAL_ELONGATION | 77 |
| ***MYC / mTOR / growth*** | |
| HALLMARK_MYC_TARGETS_V1 | 200 |
| HALLMARK_MYC_TARGETS_V2 | 58 |
| HALLMARK_MTORC1_SIGNALING | 200 |
| HALLMARK_PI3K_AKT_MTOR_SIGNALING | 105 |
| KEGG_MTOR_SIGNALING_PATHWAY | 52 |
| KEGG_PHOSPHATIDYLINOSITOL_SIGNALING_SYSTEM | 76 |
| ***Cell proliferation*** | |
| HALLMARK_E2F_TARGETS | 200 |
| HALLMARK_G2M_CHECKPOINT | 200 |
| GOBP_CELL_CYCLE | 1755 |
| GOBP_DNA_REPLICATION | 321 |
| ***DNA damage / repair*** | |
| HALLMARK_DNA_REPAIR | 150 |
| GOBP_DNA_DAMAGE_RESPONSE | 971 |
| GOBP_DNA_REPAIR | 643 |
| GOBP_HOMOLOGOUS_RECOMBINATION | 77 |
| GOBP_DOUBLE_STRAND_BREAK_REPAIR | 329 |
| ***Metabolism / stress*** | |
| HALLMARK_GLYCOLYSIS | 200 |
| HALLMARK_OXIDATIVE_PHOSPHORYLATION | 201 |
| HALLMARK_HYPOXIA | 200 |
| HALLMARK_UNFOLDED_PROTEIN_RESPONSE | 113 |
| GOBP_PROTEIN_FOLDING | 235 |
| GOBP_RESPONSE_TO_ENDOPLASMIC_RETICULUM_STRESS | 290 |
| GOBP_PURINE_DEOXYRIBONUCLEOTIDE_BIOSYNTHETIC_PROCESS | 5 |
| GOBP_PYRIMIDINE_NUCLEOTIDE_BIOSYNTHETIC_PROCESS | 34 |
| ***Residual disease phenotype*** | |
| HALLMARK_APOPTOSIS | 161 |
| HALLMARK_EPITHELIAL_MESENCHYMAL_TRANSITION | 200 |
| HALLMARK_TGF_BETA_SIGNALING | 54 |
| HALLMARK_INFLAMMATORY_RESPONSE" | 201 |

**Table S3.** Antibodies used in this study.

| Antibody | Source | Working dilution | Identifier |
|---|---|---|---|
| γH2AX | Cell Signaling Technology | 1:400 | Cat #80312S |
| γH2AX | Abcam | 1:1000 | Cat #ab2893 |
| Nucleophosmin | Abcam | 1:400 | Cat #ab10530 |
| p21 | Santa Cruz Biotechnology | 1:200 | Cat #sc-53870 |
| RSL1D1 | MilliporeSigma | 1:400 | Cat #HPA043483 |
| HCS Nuclear Mask | Life Technologies | 1:2000 | Cat #H10325 |
| Alexa 488 donkey anti-mouse | Life Technologies | 1:300 | Cat #A21202 |
| Alexa 647 donkey anti-rabbit | Life Technologies | 1:300 | Cat #A31573 |
